# CYRI1-mediated inhibition of RAC1 signalling restricts *Salmonella* Typhimurium infection

**DOI:** 10.1101/238733

**Authors:** Kyoko E Yuki, Hadir Marei, Evgenij Fiskin, Megan M Eva, Angelica A Gopal, Jeremy A Schwartzentruber, Jacek Majewski, Mathieu Cellier, Judith N Mandl, Silvia M Vidal, Danielle Malo, Ivan Dikic

**Author notes:** These authors contributed equally to this work (Kyoko E Yuki, Hadir Marei, Evgenij Fiskin, Megan M Eva). These authors jointly supervised this work (Danielle Malo, Ivan Dikic).

## Abstract

*Salmonella* presents a global public health concern. Central to *Salmonella* pathogenicity is an ability to subvert host defence mechanisms through bacterial effectors that target key host proteins implicated in restricting infection. Thus, to gain insight into host-pathogen interactions governing *Salmonella* infection, a thorough understanding of host defence mechanisms is needed. To tackle this, we performed an *in vivo* genome-wide ENU mutagenesis screen to uncover novel host defence proteins. Through this screen we identified an uncharacterised protein, which we name CYRI1 (<u>CY</u>FIP-related <u>R</u>AC1 Interacting protein <u>1</u>) that serves as a *Salmonella* resistance factor. We show that CYRI1 binds to the small GTPase RAC1 through a conserved domain present in CYFIP proteins, which are known RAC1 effectors that stimulate actin polymerisation. However, unlike CYFIP proteins, CYRI1 negatively regulates RAC1-driven actin cytoskeleton remodelling, thereby attenuating processes such as phagocytosis and cell migration. This, in turn, enables CYRI1 to counteract *Salmonella* at various stages of infection, including bacterial entry into epithelial cells, internalisation into myeloid-derived phagocytes as well as phagocyte-mediated bacterial dissemination. Together, this outlines a novel host defence mechanism that is crucial for determining bacterial fate.

---

Enteropathogenic bacteria are a major public health concern, accounting for over 300 million foodborne illnesses and 60 % of related fatalities worldwide. Among these bacteria, *Salmonella enterica* subspecies (subsp.) *enterica* serotypes are associated with the largest disease burden^1^. In fact, it is estimated that non-typhoidal *Salmonella*, alone, accounts for approximately 93.8 million illnesses and 155,000 deaths worldwide per year^2^. This, together with the increasingly widespread emergence of antibiotic-resistant *Salmonella* strains^3^, calls for a better understanding of the molecular mechanisms underlying *Salmonella* pathogenicity and the associated host cellular responses.

*Salmonella enterica* subsp. are facultative, motile, Gram-negative bacteria that invade phagocytic M cells of the Peyer’s patches (PP) as well as non-phagocytic enterocytes^4^. Upon breaching the gastrointestinal mucosa, *Salmonella* spread to the mesenteric lymph nodes (MLNs) and are engulfed by phagocytes, including macrophages, neutrophils and dendritic cells^5^. Depending on host-pathogen dynamics, phagocytes can provide a protective niche for bacterial replication^6^. Additionally, phagocyte migration can also aid dissemination to secondary tissues, such as the spleen and liver, where *Salmonella* can replicate further^4,6,7^.

Underlying *Salmonella* pathogenicity is an ability to hijack host cellular processes, through targeting key host signalling proteins^8^. This is mediated, in part, via a number of bacterial effectors that are delivered into the host cytosol using specialised molecular “syringes”, known as the Type III Secretion System (T3SS)^9,10^. Among targeted host proteins, cytoskeletal regulators are of particular importance, as their modulation by *Salmonella* effectors is crucial for mediating infection at different stages^11^. For example, *Salmonella* uptake depends, largely, on the timely activation of the host small Rho GTPases, RAC1 and CDC42. Through mimicking host Guanine-nucleotide Exchange Factors (GEFs), activators of small GTPases, bacterial effectors such as SopE and SopE2 are able to increase RAC1 and CDC42 activity^12–16^. This, triggers a number of signalling cascades within the host, including the recruitment of active RAC1 to the WAVE Regulatory Complex (WRC), a heteropentameric protein assembly comprising of CYFIP1/2, NAP1, ABI2, HSPC300 and WAVE^17,18^. Binding of active RAC1 to the WRC through CYFIP proteins induces conformational changes that alleviate complex autoinhibition leading to WRC-mediated activation of the actin nucleating ARP2/3 complex^19–21^. In turn, the ARP2/3 mediates actin polymerisation, resulting in membrane ruffling and *Salmonella* engulfment in specialised vacuoles, known as *Salmonella*-containing vacuoles (SCVs), in which *Salmonella* reside and replicate^22–24^. While in the SCVs, *Salmonella* release a second wave of effectors that target other host proteins, including additional key cytoskeletal modulators, to promote bacterial survival and dissemination^25^. Intriguingly, host cytoskeletal proteins are also implicated in antibacterial immune responses^26,27^. Thus, in addition to allowing *Salmonella* to control cellular processes, such as membrane ruffling, hijacking host cytoskeletal regulators also marks the ability of *Salmonella* to strategically target key factors implicated in host defence. As such, deciphering host defence mechanisms against *Salmonella* promises to shed light onto the different strategies employed by bacteria and to reveal novel host-pathogen interactions governing *Salmonella* infection.

To gain additional insight into the interplay between host proteins and *Salmonella* bacterial effectors during infection we set out to uncover key host defence mechanisms in an unbiased manner. We therefore performed a genome-wide recessive screen in mutagenized mice to identify host proteins implicated in conferring resistance against *Salmonella*. Through this screen we identified FAM49B, an uncharacterised protein, as a *Salmonella* resistance factor. Structural and biochemical analysis revealed that FAM49B is evolutionary related to CYFIP proteins and can similarly interact with RAC1. Based on this, we renamed FAM49B and its coding gene as CYRI1 (<u>CY</u>FIP-related <u>R</u>AC1 <u>I</u>nteracting protein <u>1</u>) to provide a more descriptive name. Intriguingly, we show that unlike CYFIP proteins, CYRI1 negatively regulates RAC1-driven actin polymerisation, which restricts bacterial entry, phagocytosis and phagocyte cell migration, thereby attenuating *Salmonella* infection. This sheds light onto the function of a previously uncharacterised negative modulator of RAC1 signalling and cytoskeletal remodelling and outlines additional strategies employed by host cells to counteract *Salmonella* infection.

## Results

### CYRI1 is a *Salmonella* resistance factor

*Salmonella enterica* subspecies *enterica* serovar Typhimurium (*S*. Typhimurium) is amongst the most predominant serotypes associated with human disease^28^. We therefore performed an *N*-ethyl-*N*-nitrosourea (ENU)-mediated genome-wide mutagenesis screen in mice aimed at identifying proteins implicated in host defence against *S*. Typhimurium. Through screening 8,389 third generation (G3) mice derived from 491 pedigrees, we identified 10 deviant pedigrees named *Immunity to Typhimurium* (*Ity*) locus 9–11 and 14–20 ^29^. Among them, G3 offspring from the *Ity15* pedigree exhibited increased susceptibility to *S*. Typhimurium infection (Fig. 1 a, b). *In vivo* imaging with bioluminescent *Salmonella* also revealed progressive bacterial load increase in *Ity15* mutants (*Ity15^m/m^*) starting as early as three days post infection (Fig. 1 c). Additionally, infected *Ity15^m/m^* mice displayed significantly higher bacterial load in the spleen and liver compared to littermate controls (Fig. 1 d). Consistently, serum pro-inflammatory (TNFα and IL-6) and anti-inflammatory (IL-10) cytokines were significantly higher in mutant mice at day 5 post-infection (supplementary Fig. 1 a). *Ity15^m/m^* mice also presented multiple abscesses in the spleen and liver, with lesions being more diffuse in the *Ity15^m/m^* spleen and the white pulp showing marked lymphocytolysis. The *Ity15^m/m^* liver also presented more parenchymal necrosis with the presence of frequent fibrin thrombi in the microvasculature when compared to controls (Supplementary Fig. 1 b). Overall, these pathological changes are indicative of the presence of an overwhelming septicaemia in *Ity15^m/m^* mice with induction of disseminated intravascular coagulation.

**Figure 1.**
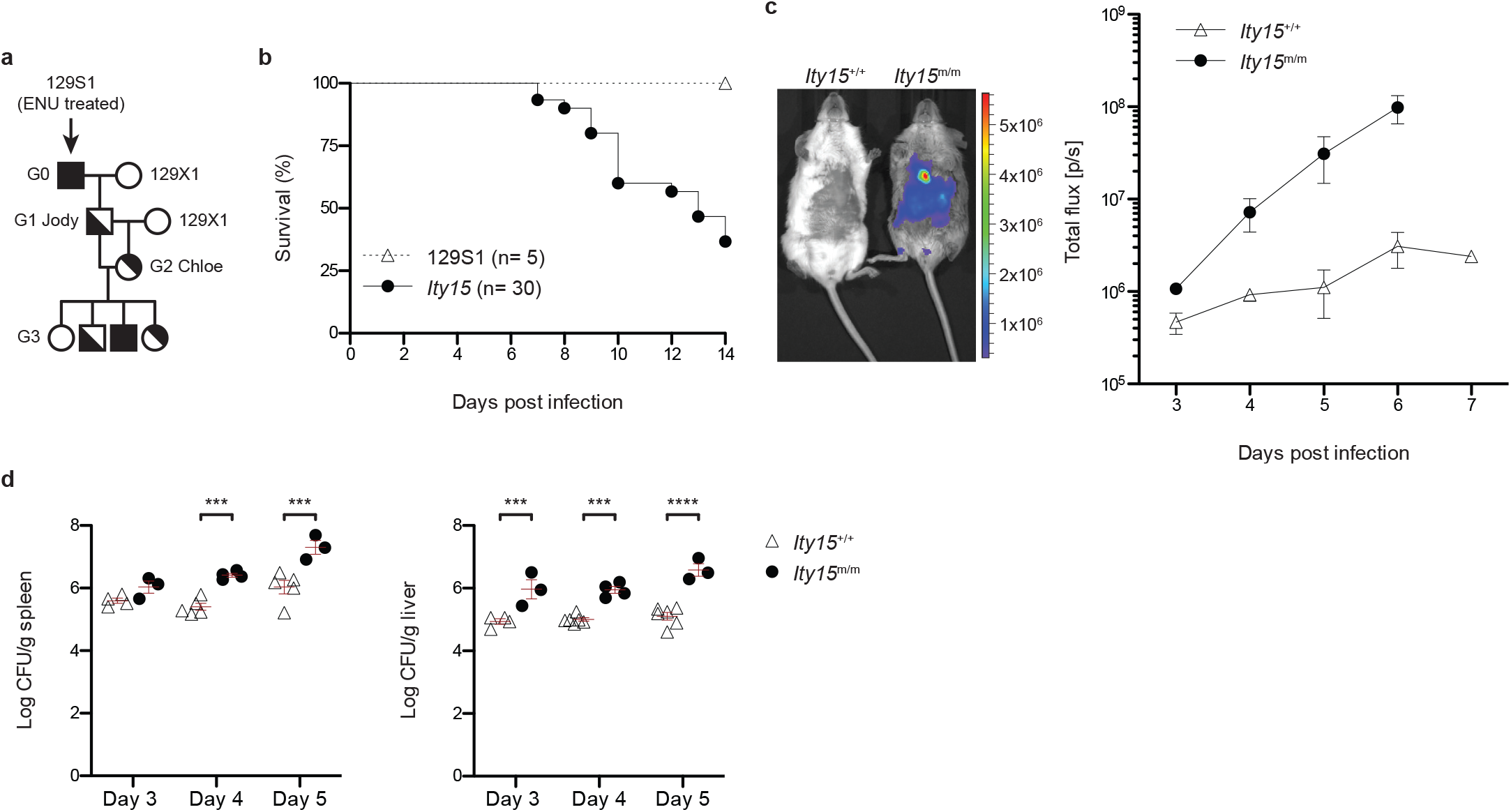
The *Ity15* pedigree displays increased susceptibility to *S*. Typhimurium infection. **a** Breeding scheme for the production of ENU-induced mutant mice; the G2 female (Chloe) was backcrossed to its G1 father (Jody) to generate the *Ity15* pedigree G3 offspring. **b** Survival curves of indicated number of G3 offspring from the *Ity15* pedigree and wild type 129S1 mice following infection with *S*. Typhimurium. Data were pooled from six independent experiments. Log-rank (Mantel-Cox) test was used to assess significance. p = 0.02. c Xen26 luminescent *S*. Typhimurium replication kinetics in *Ity15^+/+^* and *Ity15^m/m^* mice over 7 days. Graph represents the mean flux ± SEM calculated from three individual mice per genotype. Representative *Ity15^+/+^* and *Ity15^m/m^* mice are shown at day 6. **d** *Ity15^+/+^* and *Ity15^m/m^* mice spleen and liver bacterial load at day 3, 4 and 5 post infection. Graph represents log CFU count per spleen/liver weight (g) for individual mice and the mean ± SEM calculated from one experiment. Data are representative of five independent experiments. Two-way ANOVA with Bonferroni posttests was used to assess significance. ***= p ≤ 0.001; ****= p ≤ 0.0001.

To pinpoint the causative mutation underlying the *Ity15^m/m^* phenotype, we outcrossed the G1 male to DBA/2J females to generate F1 and intercrossed F1 mice to produce F2 progeny. We performed genome-wide linkage analysis in 24 F2 (9 *Salmonella*-susceptible and 15 *Salmonella*-resistant) mice using polymorphic Single Nucleotide Polymorphisms (SNPs) between 129S1 and DBA/2J. This revealed a single linkage peak on chromosome 15 with an LOD score of 4.54 between markers rs3677062 and rs13482628, delineating an interval of 4.4 Mb (Supplementary Fig. 2 a). F2 mice homozygous for the 129S1 mutant allele (*Ity15^m/m^*) at the peak marker on chromosome 15 succumbed to infection by day 8 compared to heterozygous (*Ity15+^m^*) and wild type (*Ity15^+/+^*) mice (Supplementary Fig. 2 b). Interestingly, exome sequencing performed in two susceptible F2 animals identified four ENU-induced homozygous Single Nucleotide Variants (SNVs). Importantly, one of the identified SNVs was found in *Cyri1* (*Fam49b*), an uncharacterised protein-coding gene, that maps to the critical interval on chromosome 15. The mutation within *Cyri1* consisted of a T to A substitution within the splice donor site of exon 9 resulting in exon 9 skipping and causing a frameshift and a premature stop codon within exon 10 (p.N212GfsX9) (Fig. 2 a and Supplementary Fig. 2 c). Indeed, biochemical analysis of spleen and liver tissues revealed that the splicing mutation abrogates CYRI1 protein expression in *Ity15^m/m^* mice (Fig. 2 b).

**Figure 2.**
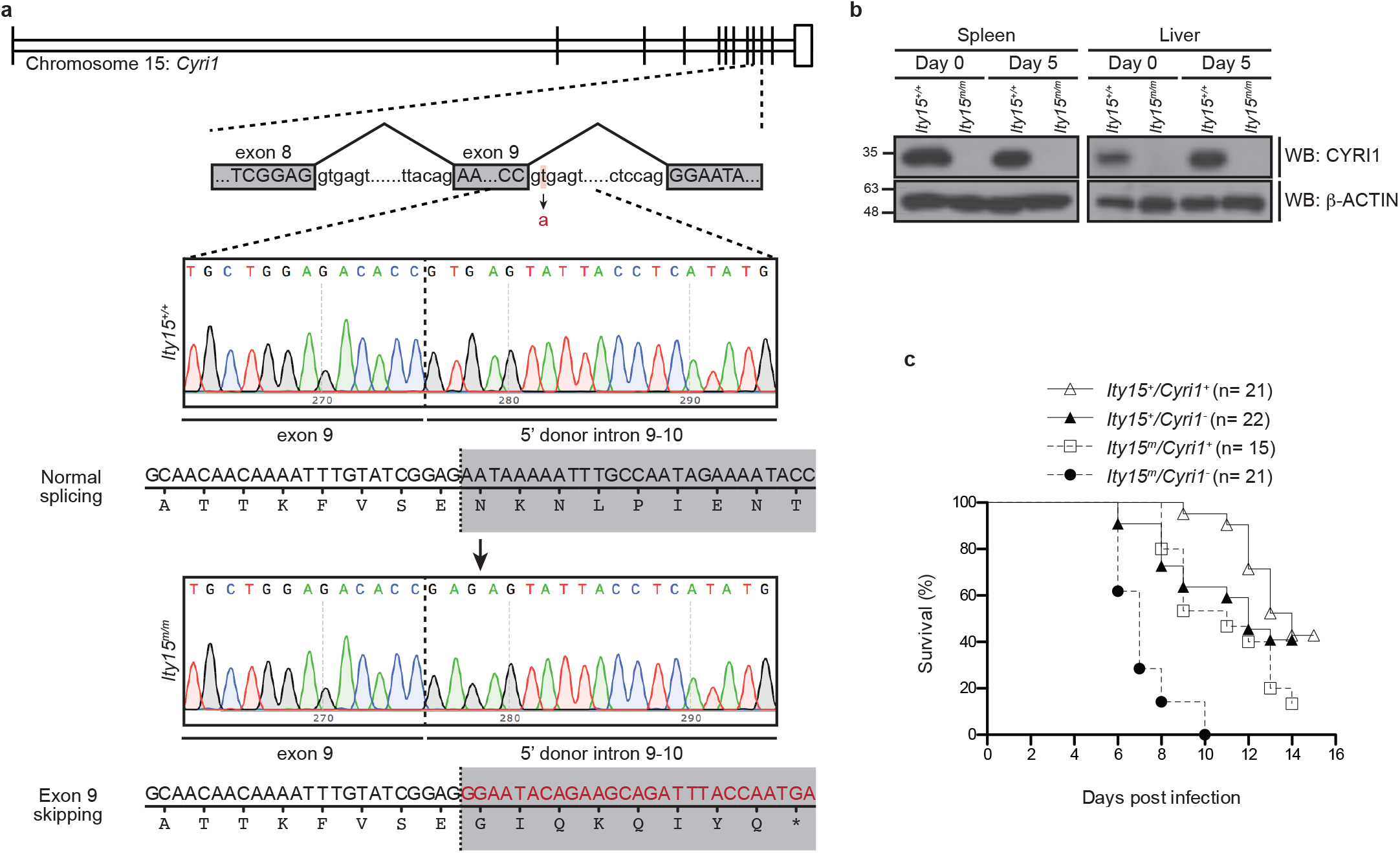
A mutation in *Cyri1* confers susceptibility to *S*. Typhimurium infection in the *Ity15* pedigree. **a** Schematic representation of the ENU-induced *Cyri1* mutation. **b** Western blot analysis of CYRI1 protein expression in *Ity15^+/+^* and *Ity15^m/m^* mice spleen and liver at day 0 and 5 post infection, representative of three independent experiments. β-ACTIN was used as a loading control. **c** Allelic complementation assay. Survival curves of indicated number of F1 mice derived from a cross between *Ity15^+/m^* and *Cyri1^+/−^*. Data were pooled from four independent experiments. Log Rank (Mantel-Cox) test was used to assess significance. p < 0.0001.

To further validate *Cyri1* as the causative gene underlying susceptibility in *Ity15^m/m^* mice, we performed allelic complementation in which heterozygous *(Ity15^+/m^)* mice were crossed to mice heterozygous for a knockout (KO) allele at *Cyri1* (*Cyri1^+/−^*). We next assessed survival in F1 animals upon *S*. Typhimurium infection to ascertain susceptibility (Supplementary Fig. 2 d). Interestingly, we observed a lack of complementation with *Ity15^m^/Cyri1^−^* mice being as susceptible as *Ity15^m/m^* mice (Fig. 2c), confirming that the observed *Cyri1* mutation is responsible for *Salmonella* susceptibility in *Ity15^m/m^* mice. This identifies *Cyri1* as a *Salmonella* resistance gene with loss of CYRI1 expression resulting in enhanced *Salmonella* dissemination and septic shock.

### CYRI1 is a CYFIP-related RAC1 effector

Given the uncharacterised nature of CYRI1, we next explored its potential cellular functions. Primary structural analysis revealed that CYRI1 shares 80 % sequence identity with FAM49A, a closely related uncharacterised protein, within a single domain, Domain of Unknown Function 1394 (DUF1394), which is found in two other proteins in the human proteome: CYFIP1 and CYFIP2 (Supplementary Fig. 3 a).

CYFIP proteins are part of the WRC and are integral in relaying information from active RAC1 to the ARP2/3 complex, by directly binding to RAC1^19^. This raised the possibility that CYRI1 might also bind to active RAC1. Indeed, *in vitro* biochemical assays demonstrated that CYRI1 associates with RAC1 (Fig. 3 a). The CYRI1-RAC1 interaction was also observed in cells, as evident by duolink *in situ* proximity ligation assays using antibodies against endogenous RAC1 and endogenous or exogenous CYRI1 (Fig. 3 b, c). Moreover, GST pulldown experiments using purified GST-RAC1 loaded with either GDP or GTPγS to mimic inactive and active RAC1, respectively, revealed that CYRI1 binds preferentially to active RAC1 (Fig. 3 d). Consistently, endogenous RAC1 from HEK293T lysates pre-treated with GTPγS displayed enhanced binding to GST-CYRI1 compared to endogenous RAC1 from GDP pretreated lysates (Fig. 3 e, f).

**Figure 3.**
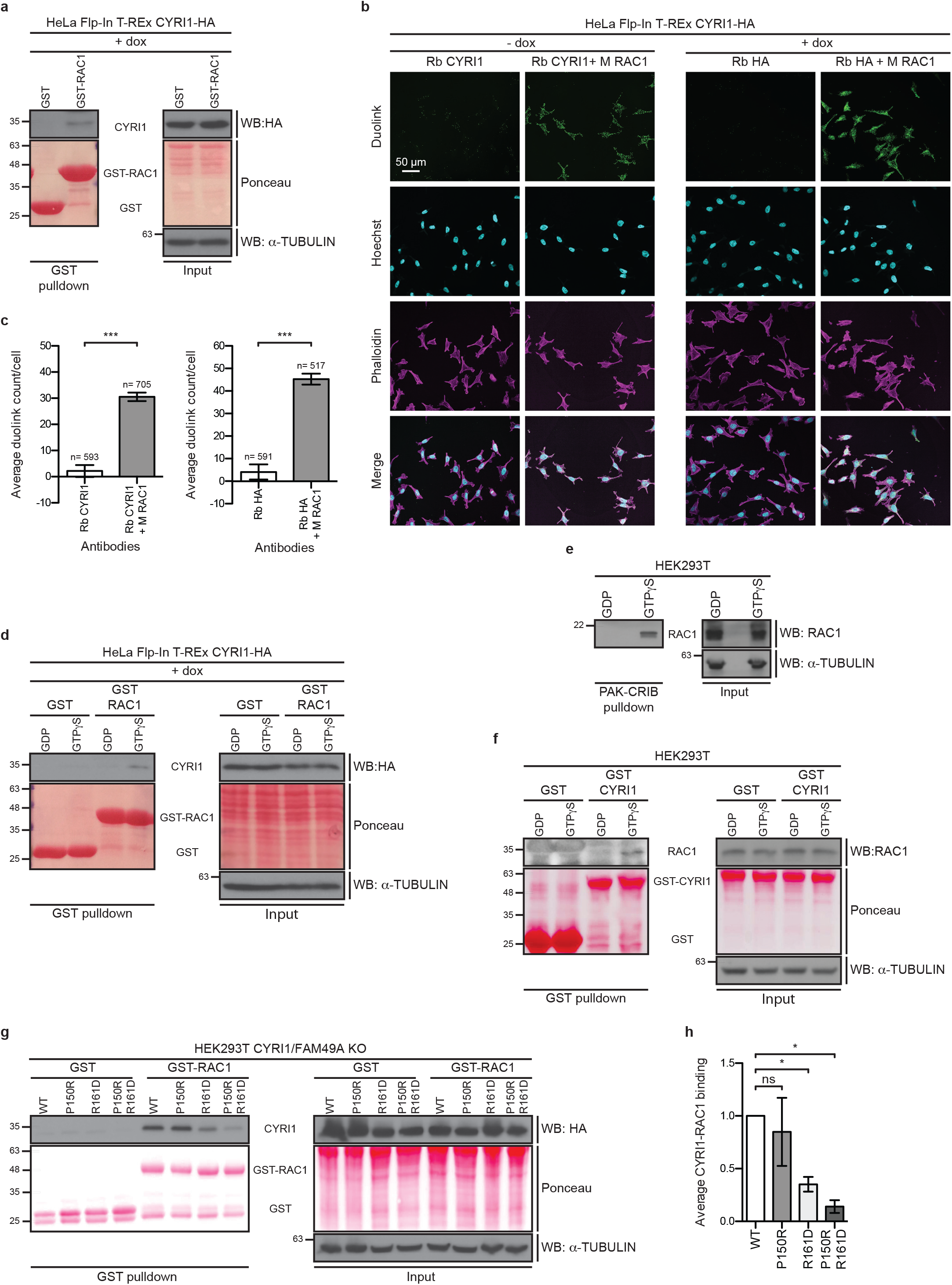
CYRI1 binds to the small GTPase RAC1 in a GTP-dependent manner. **a** GST pulldown of GST or GST-RAC1 incubated with lysates derived from doxycycline (dox)-treated HeLa Flp-In T-REx CYRI1-HA cells. Co-precipitated CYRI1-HA was detected by western blotting. **b** Duolink *in situ* PLA immunofluorescence images of untreated (-dox) or dox-treated (+ dox) HeLa Flp-In T-REx CYRI1-HA cells using indicated antibodies, representative of three independent experiments. Phalloidin and Hoechst were used to visualise the actin cytoskeleton and nuclei, respectively. **c** Quantification of average duolink count per cell from indicated number of cells ± SD calculated from one biological replicate out of three independent experiments. Unpaired two-tailed t-test with Welch’s correction was used to assess significance. ***= p ≤ 0.001. **d** GST pulldown of GDP or GTPγS-loaded GST or GST-RAC1 incubated with lysates derived from dox-treated HeLa Flp-In T-REx CYRI1-HA cells. Co-precipitated CYRI1-HA was detected by western blotting. **e** PAK-CRIB pulldown of GDP or GTPγS-treated HEK293T lysates from one biological replicate. Active and total RAC1 were detected by western blotting. α-TUBULIN was used as a loading control. f GST pulldown of GST or GST-CYRI1 incubated with GDP or GTPγS-treated HEK293T lysates from one biological replicate. Co-precipitated RAC1 was detected by western blotting. **g** GST pulldown of GTPγS-loaded GST or GST-RAC1 incubated with lysates derived from HEK293T CYRI1/FAM49A knockout (KO) cells expressing indicated CYRI1 mutants. Coprecipitated CYRI1-HA was detected by western blotting. For a, d, f and g, α-TUBULIN and ponceau were used as loading controls. For a, d and g panels are representative of three independent experiments. **h** Quantification of CYRI1 binding to GST-RAC1. Graph represents average CYRI1-RAC1 binding ± SEM normalised to CYRI1-HA wild type (WT)-RAC1 binding calculated from three independent experiments. One-way ANOVA with Dunnett’s multiple comparison test was used to assess significance relative to CYRI1 WT. ns= non-significant; *= p ≤ 0.05.

Despite a weak sequence homology of ~ 20 %, the HHpred^30^ predicted CYRI1 structure suggests that CYRI1 shares significant structural similarity with the CYFIP1 DUF1394 domain (score= 493.83, e-value=1.5e-58) (Supplementary Fig. 3 b). A reported WRC structure indicated the presence of two RAC1 binding sites, cysteine 179 (C179) and arginine 190 (R190), within the CYFIP1 DUF1394 domain, which abrogate RAC1 binding when mutated to R and aspartic acid (D), respectively^21^. In CYRI1, proline 150 (P150) and R161 correspond to CYFIP1 C179 and R190, respectively (Supplementary Fig. 3 c, d). We therefore generated P150R and R161D single and double CYRI1 mutants to assess their RAC1 binding ability. To eliminate any potential non-specific binding due to dimerization between the mutants and endogenous CYRI1 or FAM49A, we expressed these mutants in a CYRI1/FAM49A double KO HEK293T cell line (HEK293T CYRI1/FAM49A KO) and performed GST pulldown using GTPγS-loaded purified GST-RAC1. Consistent with the high conservation between CYRI1 R161 and CYFIP1 R190, the R161D mutation significantly reduced RAC1 binding. In contrast, the CYRI1 P150R mutation had no effect on CYRI1-RAC1 interaction (Fig. 3 g, h). However, the P150R/R161D CYRI1 double mutant displayed a greater reduction in RAC1 binding compared to the R161D mutant alone, suggesting that while non essential, P150 also contributes to RAC1 interaction (Fig. 3 g, h). Together, this demonstrates that CYRI1 is evolutionary related to CYFIP proteins and can similarly interact with RAC1.

### CYRI1 negatively regulates RAC1 signalling

Both the structural homology to CYFIP proteins and its ability to bind to RAC1 hinted at a potential role of CYRI1 in regulating RAC1-driven actin cytoskeletal modulation. To test this, we generated CYRI1 KO HeLa cells and assessed their morphology (Fig. 4 a). CYRI1 KO cells exhibited a round, unpolarised and flattened pancake-like morphology with extensive membrane ruffling and increased cellular spread and circularity compared to parental or control KO HeLa cells (Fig. 4 b-e). These morphological changes were also accompanied by extensive actin cytoskeletal rearrangements, as indicated by phalloidin, ACTIN and RAC1 immunofluorescence staining (Fig. 4 f and Supplementary Fig. 4 a).

**Figure 4.**
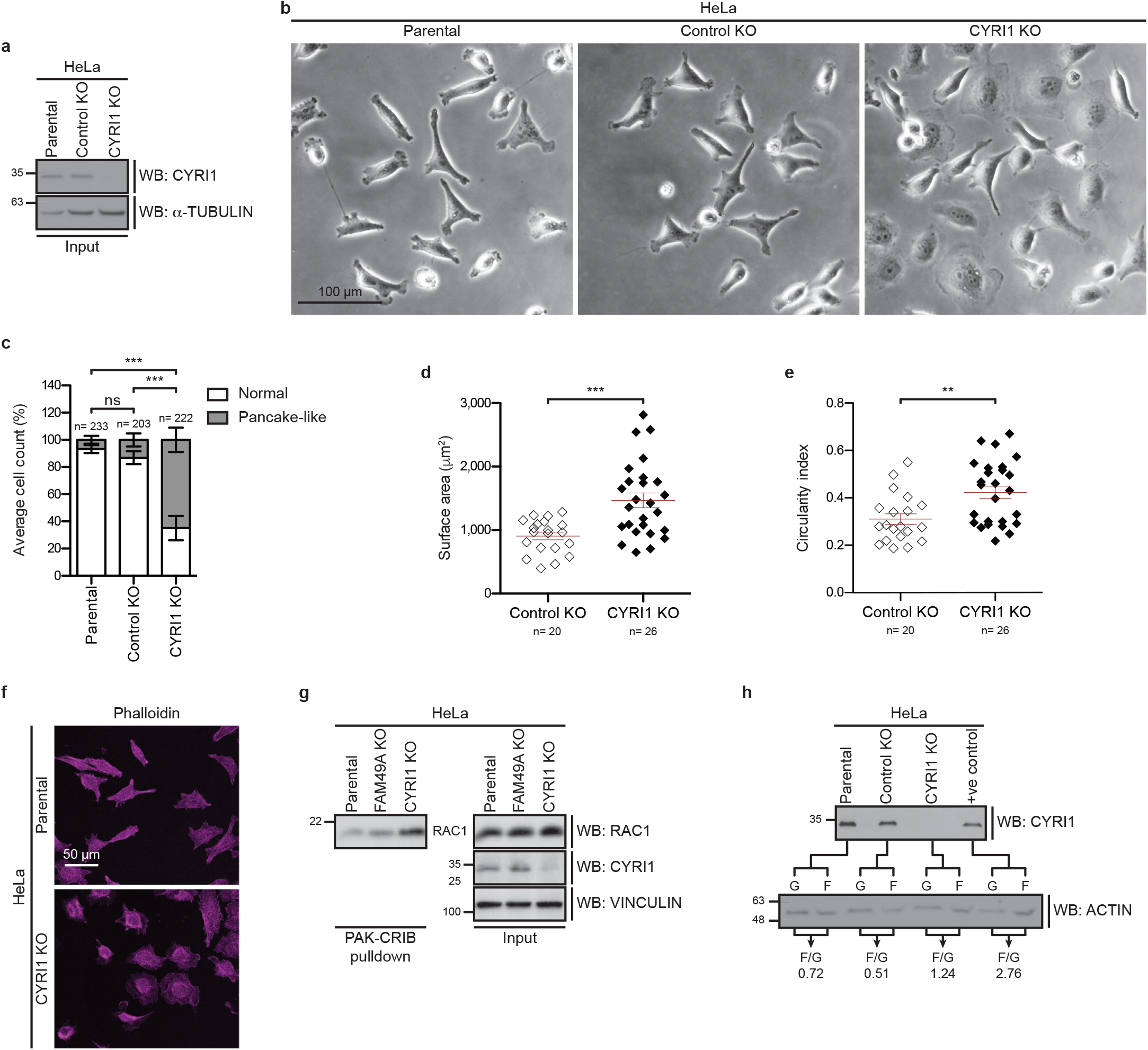
CYRI1 regulates cell morphology and inhibits RAC1-driven actin polymerisation. **a** Western blot analysis of lysates from parental, control or CYRI1 knockout (KO) HeLa cells to determine the efficiency of CYRI1 KO, representative of three independent experiments. α-TUBULIN was used as a loading control. **b** Phase-contrast images of parental, control KO or CYRI1 KO, representative of three independent experiments. **c** Quantification of cellular phenotypes outlined in (b). Graph represents % of cells with outlined morphology categories ± SEM from indicated number of cells per condition calculated from three independent experiments. Two-way ANOVA with Bonferroni posttests was used to assess significance. ns= non-significant; ***= p ≤ 0.001. **d, e** Quantification of cell area and shape parameters from indicated number of control KO and CYRI1 KO HeLa cells. Graph represents values of surface area (d) and circularity index (e) for individual cells and the mean ± SEM calculated from three independent experiments. Unpaired two-tailed t-test was used to assess significance. **= p ≤ 0.01; ***= p ≤ 0.001. **f** Immunofluorescence images of parental and CYRI1 KO cells using phalloidin to visualise the actin cytoskeleton, representative of two independent experiments. **g** PAK-CRIB pulldown of lysates from control KO, FAM49A KO or CYRI1 KO HeLa cells from one biological replicate. Active and total RAC1 were detected by western blotting. α-TUBULIN was used as a loading control. **h** G/F actin *in vivo* assay from parental, control KO, CYRI1 KO HeLa cells and HeLa cells treated with an actin-polymerising agent as a positive control (+ve control) from one biological replicate. G-actin, F-actin and CYRI1 protein levels were detected by western blotting.

Intriguingly, HeLa cells expressing the RAC1 activating mutant, RAC1 P29S displayed a round and flattened morphology similar to that observed upon CYRI1 deletion (Supplementary Fig. 4 b-d), which is consistent with previous observations linking the pancake-like morphology to RAC1 hyperactivation^31^. This suggested that the morphological changes associated with CYRI1 KO might be due to increased RAC1 activation, leading to intensified cytoskeletal dynamics. Indeed, increased active RAC1 levels were observed in CYRI1 KO cells (Fig. 4 g). Additionally, CYRI1 KO HeLa cells displayed enhanced actin polymerisation, as measured by an increased filamentous (F)-actin to monomeric (G)-actin ratio when compared to parental or control KO HeLa cells (Fig. 4 h). Together, this identifies CYRI1 as a negative regulator of RAC1-driven actin cytoskeletal remodelling.

### CYRI1 restricts *Salmonella* entry and dissemination by modulating cytoskeletal processes

RAC1 activation and subsequent actin polymerisation is central for *Salmonella* internalisation into non-phagocytic enterocytes^24^. We therefore hypothesised that CYRI1 might impede bacterial entry. To investigate this, we assessed bacterial entry in parental, control KO or CYRI1 KO HeLa cells following *Salmonella* infection as a model of bacterial internalisation into non-phagocytic enterocytes in the gut. Indeed, CYRI1 deletion resulted in increased *Salmonella* internalisation relative to parental and control KO cells (Supplementary Fig. 5 a, b), indicating that CYRI1 inhibits actin cytoskeletal rearrangements required for bacterial entry.

RAC1-driven actin cytoskeleton remodelling also plays a major role in mediating phagocytosis^32^ and cell motility^33^. This is particularly relevant in myeloid-derived phagocytes, such as macrophages and neutrophils, which can foster bacterial replication and dissemination *in vivo*^4,6,7,34,35^. We therefore evaluated the role of CYRI1 in cells of haematopoietic origin for conferring *Salmonella* resistance by generating bone marrow chimeric mice. Strikingly, the transfer of *Ity15^m/m^* mice bone marrow cells into wild type mice resulted in increased *Salmonella* susceptibility as evident by reduced survival upon infection. Conversely, increased resistance could be conferred to *Ity15^m/m^* mice by transferring wild type bone marrow cells into mutant mice (Fig. 5 a). To further delineate the role of CYRI1 within the haematopoietic compartment, we next generated myeloid-specific conditional CYRI1 KO mice (*Cyri1^−/flox^*) (Supplementary Fig. 5 c). Loss of CYRI1 expression in myeloid cells was associated with a significant decrease in survival upon *S*. Typhimurium infection (Fig. 5 b). We also observed an increase in several serum pro-inflammatory cytokines and chemokines produced by activated neutrophils and monocytes (Supplementary Fig. 5 d and Supplementary Data Table 1) and higher liver bacterial load in *Cyri1^−/flox^* mice compared to their wild type counterparts upon *S*. Typhimurium infection (Fig. 5 c). Together, this demonstrates the importance of CYRI1 within the haematopoietic compartment for conferring resistance to *Salmonella* infection *in vivo*.

**Figure 5.**
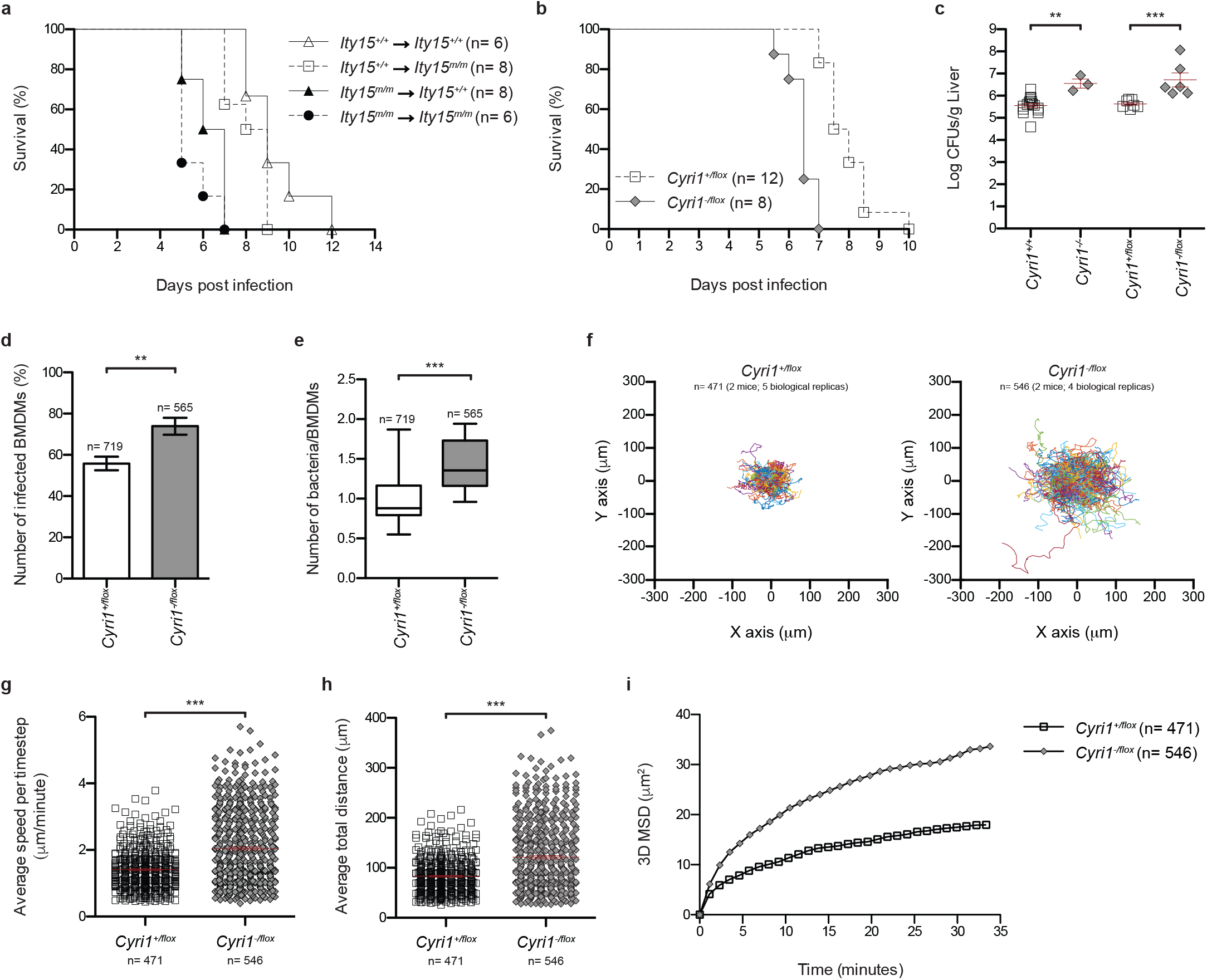
CYRI1 expression in haematopoietic cells confers resistance against *S*. Typhimurium infection. **a** Survival curves of indicated number of bone marrow chimeras in which bone marrow cells were transferred between mice as indicated by arrows, representative of two independent experiments. Log-rank (Mantel-Cox) test was used to assess significance. p= 0.0012. **b** Survival curves of wild type (*Cyri1*^+/flox^) and myeloid-specific CYRI1 knockout (KO) (*Cyri1*^−/flox^) mice, representative of two independent experiments. Log-rank (Mantel-Cox) test was used to assess significance. p= 0.0015. **c** Liver bacterial load from *Cyri1* wild type (*Cyri1*^+/+^), *Cyri1* full KO (*Cyri1*^−/−^), *Cyri1*^−/flox^ and *Cyri1*^−/flox^ mice 4 days after infection. Graph represents log CFU count per liver weight (g) for individual mice and the mean ± SEM calculated from one experiment. Data are representative of two experiments. One-way ANOVA with Dunnett’s multiple comparisons test was used to assess significance. ** = p<0.01; *** = p<0.001. **d** Quantification of number of infected BMDMs derived from *Cyri1*^+/flox^ and *Cyri1*^−/flox^ mice. Graph represents % of cells infected per condition ± SEM from indicated number of cells calculated from three independent experiments. **e** Quantification of number of bacteria per BMDMs derived from *Cyri1*^+/flox^ and *Cyri1*^−/flox^ mice. Graph represents the mean ± SEM calculated from three independent experiments. For d and e unpaired two-tailed t-test was used to assess significance. **= p ≤ 0.01; ***= p ≤ 0.001. **f** Normalised tracks of *Cyri1*^+/flox^ and *Cyri1*^−/flox^-derived neutrophils migrating for 1 hour. g Average speed per time-step from indicated number of *Cyri1*^+/flox^ and *Cyri1*^−/flox^-derived neutrophils. Graph represents values for individual cells and the mean ± SEM. **h** Average total distance travelled from indicated number of *Cyri1*^+/flox^ and *Cyri1*^−/flox^-derived neutrophils. Graph represents values for individual cells and the mean ± SEM. For g and h unpaired two-tailed t-test with Welch’s correction was used to assess significance. ***= p ≤ 0.001. i 3D Mean Squared Displacement (MSD) plots of indicated number of *Cyri1*^+/flox^ and *Cyri1*^−/flox^-derived neutrophils. For f-i, data were pooled from five (*Cyri1*^+/flox^) or four (*Cyri1*^−/flox^) independent experiments.

To interrogate whether the increased *Salmonella* susceptibility observed upon CYRI1 KO in myeloid cells is due to deregulation of RAC1-driven cellular effects we analysed the phagocytic potential of Bone Marrow-Derived Macrophages (BMDMs) from control and *Cyri1^−flox^* mice. Interestingly, the percentage of *Salmonella-infected* BMDMs was significantly higher in *Cyri1^−/flox^*-derived BMDMs compared to control BMDMs (Fig. 5 d). Additionally, the percentage of BMDMs harbouring more than 1 *Salmonella* per cell was greater in *Cyri1^−/flox^*-derived BMDMs (Fig. 5 e and Supplementary Fig. 5 e), underlining a role of CYRI1 in controlling macrophage phagocytic potential, thereby restricting bacterial infection.

In addition to utilising phagocytes as a replicative niche, *Salmonella* also stimulate phagocyte migration to aid dissemination to secondary tissues, such as the spleen and liver^4,6,7^. RAC1 is known to guide neutrophil trafficking between the bone marrow and the blood as well as cell migration within tissues^36^. To further investigate whether the phenotype observed in infected *Cyri1^−/flox^* mice is due to deregulation of RAC1 signalling, we also examined the impact of CYRI1 deletion on neutrophil motility in a 3D collagen matrix as a model of their movement *in vivo*. CYRI1 deletion significantly affected the migratory behaviour of neutrophils (Fig. 5 f). Strikingly, CYRI1 KO cells displayed higher velocities (Fig. 5 g) and travelled greater distances than control cells in all three dimensions (Fig. 5 h, i). CYRI1 KO neutrophils also had significantly reduced turning angles (Supplementary Fig. 5 f) such that their directional persistence was reduced, suggesting that neutrophils were switching to a more random mode of migration in the absence of CYRI1, which is consistent with RAC1 hyperactivation^37^ These changes in the regulation of neutrophil migration upon CYRI1 KO could help explain the enhanced *Salmonella* dissemination observed in *Cyri1^−/flox^* mice, thus implicating CYRI1 in restricting this process.

## Discussion

Bacterial pathogens and host cells are in a constant “arms race” for survival. While *Salmonella* evolved various means to hijack key host proteins implicated in defence against infection^38^, host cells also devised multiple mechanisms to combat infection^39^. Given the high disease burden associated with *Salmonella* infection^1^ and the complexity of host-pathogen interactions governing *Salmonella* pathogenicity, it is crucial to gain a better understanding of the different strategies employed by pathogens as well as host cells in the course of an infection. In an attempt to tackle this, we performed an *in vivo* genome-wide ENU mutagenesis recessive screen aimed at identifying novel host *Salmonella* resistance proteins. Here, we describe the role of CYRI1, a novel cytoskeletal regulator, in counteracting *Salmonella* infection. In particular, we show that through binding to the small GTPase RAC1 and negatively regulating its activity and downstream actin polymerisation, CYRI1 restricts *Salmonella* at multiple stages of infection. This highlights an intricate mechanism by which one host protein is able to modulate host resistance through targeting *Salmonella* at critical stages of its infection cycle, such as bacterial entry to different cell types and bacterial dissemination (Fig. 6).

**Figure 6.**
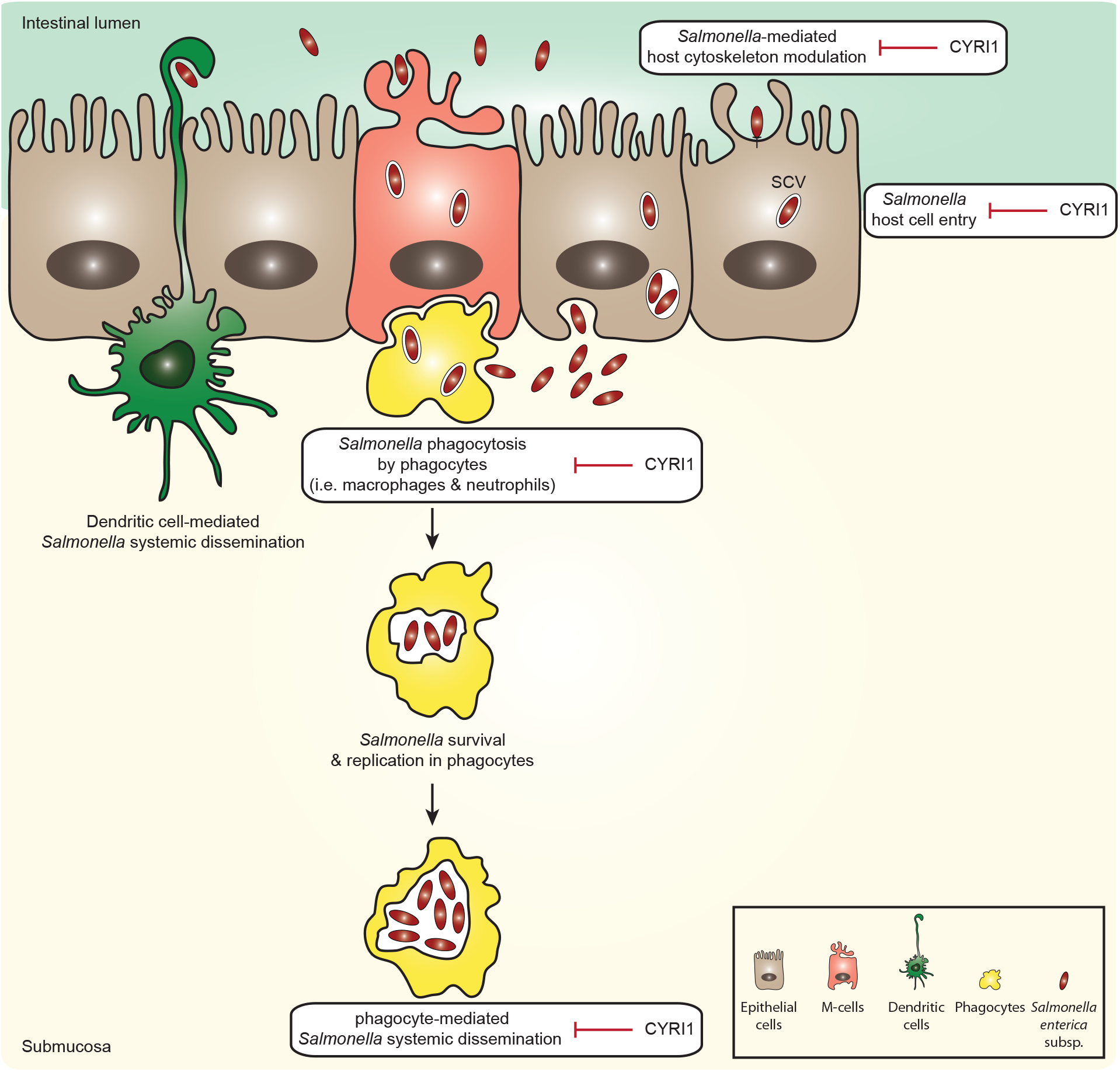
Model of CYRI1 *anti-Salmonella* effects. CYRI1 serves as a *Salmonella* resistance factor within the host through binding to RAC1 and negatively regulating RAC1-driven signalling and downstream actin polymerisation. This mechanism counteracts *Salmonella* at various stages during infection, such as impeding bacterial entry into epithelial cells, restricting bacterial phagocytosis and obstructing dissemination through negatively regulating phagocyte cell migration.

CYRI1 is a previously uncharacterised highly conserved protein, which according to the human protein atlas^40^ is ubiquitously expressed. Using *in silico* structural analysis, we show that CYRI1 and FAM49A are evolutionary related to CYFIP proteins. Of particular interest was the observation that while harbouring very little sequence homology overall, CYRI1 and CYFIP proteins are highly homologous within the DUF1394 domain, both on a primary and tertiary structural level. Importantly, our biochemical data also demonstrate a similar RAC1 binding platform within CYRI1 and CYFIP proteins, identifying the DUF1394 domain as a putative RAC1 binding motif.

RAC1-downstream signalling is implicated in both promoting and inhibiting bacterial infection^41^. Here, we demonstrate that modulation of cellular processes dependent on RAC1-driven actin cytoskeleton remodelling in the haematopoietic compartment is crucial for determining bacterial fate. We show that loss of CYRI1 leads to increased bacterial phagocytosis in macrophages and enhanced cellular migration in neutrophils. Increased levels of tissue macrophages, neutrophils and dendritic cells are a hallmark of *Salmonella* infection^42^. In fact, macrophages are amongst the most highly enriched phagocytes in PP and MLNs, followed by neutrophils during oral *Salmonella* infection^5^. Thus, once across the gastrointestinal mucosa, *Salmonella* are engulfed by macrophages and neutrophils, where they reside within the SCV. This aids bacterial replication and evasion of host innate immune response^34^ as well as bacterial dissemination and systemic infection^6^. Thus, our data implicates CYRI1 in counteracting systemic infection via restricting *Salmonella* phagocytosis and limiting bacterial dissemination, through negatively regulating RAC1 signalling.

Remodelling of the host cytoskeletal network is a key feature of many enteropathogenic bacteria. As such, cytoskeletal regulators are amongst the most commonly exploited host proteins by pathogenic bacteria^16,24^. This raises the possibility that *Salmonella* might regulate CYRI1 cellular functions upon infection, particularly given its role in counteracting Salmonella-mediated cytoskeletal rearrangements. Indeed, evidence from the literature seems to suggest that CYRI1 is a *Salmonella* targeted host protein. For instance, it has been shown that CYRI1 mRNA levels are differentially reduced upon *Salmonella* infection^43^. Additionally, using quantitative mass spectrometry, we recently identified global host ubiquitinome changes following *S*. Typhimurium infection, highlighting the ability of *Salmonella* to modulate host signalling cascades via regulating ubiquitylation of host proteins. Interestingly, CYRI1 was among the identified proteins and displayed increased ubiquitylation on lysine (K) 74 upon proteasomal inhibition, suggesting that CYRI1 is targeted for proteasomal degradation upon infection^44^. This, together with our *in* vivo infection data help outline a novel host-pathogen signalling axis involving CYRI1 that is crucial for determining bacterial fate.

In addition to bacterial infection, deregulation of RAC1 signalling is detrimental for normal cellular functioning and is implicated in a wide range of human diseases, including cancer, cardiovascular diseases and neurodegenerative disorders^41^. Thus, identifying CYRI1 as a RAC1 signalling inhibitor is likely to play a role in cellular contexts beyond bacterial infection. This calls for additional studies in the future to examine the role of CYRI1 under physiological and other pathological conditions.

## Methods

### Antibodies

Antibodies utilised in this study are outlined in Supplementary Methods Table 1.

### Mice

All animal experiments were performed under guidelines specified by the Canadian Council for Animal Care. The animal-use protocol was approved by the McGill University Animal Care Committee (protocol no. 5797). 129S1, 129X1, C57BL/6J, DBA/2, B6.129S4-*Meox2^tm1(cre)Sor^/J* (Meox-Cre, JAX stock #003755) and B6.129P2-Lyz2^tm1(cre)Ifo^/J (Lysm-Cre, JAX stock #004781) mice were purchased from the Jackson Laboratory (Bar Harbor, ME, USA). C57BL/6N-*Fam49b*^tm1a(KOMP)Wtsi^/Tcp (*Fam49b*^tm1a^) mice were obtained from the NorCOMM (North American Conditional Mouse Mutagenesis) Project. Flp mice were a generous gift from Yojiro Yamanaka (McGill Cancer Research Centre, Montreal, Canada).

### ENU mutagenesis

Generation 0 (G0) 129S1 males were mutagenized with a single intraperitoneal injection of 150 mg per kg of body weight of *N*-ethyl-*N*-nitrosourea (ENU). G0 males were outcrossed to 129X1 females to create G1 progeny that were subsequently outcrossed to 129X1 females. Resulting G2 females were backcrossed to parental G1 males to produce the G3 mice that were assessed for susceptibility to *Salmonella* infection. The G1 male from the *Ity15* pedigree was outcrossed to a DBA/2J female to generate F1 offspring. F1 mice were intercrossed to generate F2 offspring, which were then used for mapping. Crossing *Ity15^+/m^* with *Cyri1^+/−^* generated mice used for the allelic complementation assay. The *Ity15^m/m^* allele was then transferred to 129X1 background by serial backcrossing (n= 9).

### *In vivo Salmonella* infection and bioluminescence live imaging

Mice were injected intravenously in the caudal vein with 5,000 Colony Forming Units (CFUs) of *S*. Typhimurium (strain Keller). Mice were euthanized on days 3, 4 or 5 post infection to harvest spleen and liver as previously described^45^. For whole-body imaging, mice were injected intravenously with 8.8 × 10^4^ CFUs of *S*. Typhimurium strain XEN26 (PerkinElmer, 119230), anesthetized with isoflurane, anteriorly shaved, and imaged daily in an IVIS Spectrum system (PerkinElmer, 124262) until they reached clinical endpoints, at which time they were euthanized. Bioluminescent images were acquired with an open emission filter, binning factor of 16 and exposure time ranging from 2–5 minutes. The regions of interest were selected, normalised across time points, and quantified using Living Image software v.4.3.1 (PerkinElmer).

### Histology

Tissues were collected and fixed in 10 % neutral buffered formalin for 24 hours at 20 °C, then placed in 70 % ethanol at 4 °C before processing and embedding (Goodman Cancer Research Center histology facility, McGill University). Embedded tissues were sectioned and stained with hematoxylin and eosin.

### Cytokine analysis

Blood was collected from uninfected and infected *Ity15^+/+^, Ity15^m/m^, Cyri1^+/flox^ and Cyri1^−/flox^* by cardiac puncture into serum separating tubes (Sarstedt). Serum cytokines were determined either using ELISA kits (TNFα, IL-6 and IL-10) purchased from eBiosciences (Ready-SET-Go! kits) or using mouse cytokine/chemokine 32-multiplex Luminex array (Eve Technologies, Calgary, Canada).

### Genetic mapping, genotyping and exome sequencing

DNA was extracted from a tail biopsy by proteinase K digestion and phenol-chloroform extraction. The genome scan was performed using the Medium Density SNP Panel from Illumina (708 SNPs were informative between 129S1 and DBA/2 strains) (The Centre for Applied Genomics, Toronto, Canada). The mapping was done using 24 mice (9 *Salmonella*-susceptible and 15 *Salmonella*-resistant). Exome sequencing was done in two susceptible mice. Briefly, exon capture was performed with the SureSelect Mouse All Exon Kit (Agilent Technologies, 5190-4641) and sequencing of 100 bp paired end reads on Illumina HiSEquation 2000 generating over 8 Gb of sequence for each sample (Centre National de Génotypage, Évry, France). Read alignment, variant calling and annotation were performed as previously described^46^. The *Ity15* mutation in *Cyri1* was subsequently genotyped using a Custom Taqman SNP genotyping assay (Applied Biosystems).

### *In silico* analysis of CYRI1

Uniprot was used to obtain human protein sequences for CYRI1/FAM49B (Q9NUQ9), FAM49A (Q9H0Q0), CYFIP1 (Q7L576) and CYFIP2 (Q96F07). Schematic representation of domains present in these proteins was based on the Pfam database (http://pfam.xfam.org). The CYRI1 structure and its homology to CYFIP1 were predicted using the HHpred software^30^. Structural models presented in the manuscript were generated in MacPymol based on HHpred structural predictions. The presence of CYRI1, FAM49A, CYFIP1 and CYFIP2 in other species was determined using a Uniprot BLAST search (http://www.uniprot.org/blast). Important model organisms containing the respective proteins were used for sequence alignment, focusing on a region surrounding known RAC1-binding residues within the CYFIP1 DUF1394 domain. Sequence alignment and evolutionary conservation across the indicated species was conducted using Jalview.

### Cell culture

All cell lines were cultured in Dulbecco’s Modified Eagle Medium (DMEM; Thermo Fisher Scientific, 11960-044) supplemented with 10 % v/v fetal bovine serum (FBS; Thermo Fisher Scientific; 10270106) and 10 μg/ml penicillin-streptomycin (Sigma Aldrich, P0781-100 ml) and kept at 37 °C and 5 % CO_2_.

### Constructs

Details of plasmids used in this study are summarised in Supplementary Methods Table 2.

### Generation of cell lines

Transfection with indicated constructs was performed using GeneJuice (Nalgene, 16211-034) according to manufacturer’s instructions. For overexpression, CYRI1-HA was sub-cloned into pcDNA5/FRT/TO and the doxycycline (dox)-inducible cell lines were generated using the Flp-In T-REx system (Thermo Fisher Scientific, K650001) according to manufacturer’s instructions. HeLa cells containing the dox-inducible expression system were subjected to antibiotic selection with blasticidin (15 mg/ml; Capricorn Scientific; BLA-5x), zeocin (1 mg/ml; Invitogen, 45-0430) and hygromycin B (250 μg/ml; Invitrogen, 10687010). Retroviral transduction was performed as previously described^47^. For CRISPR/Cas9-medaited knockout (KO), three sgRNAs, outlined in Supplementary Methods Table 3, were selected from The Broad Institute Genetic Perturbation Platform (https://portals.broadinstitute.org/gpp/public/analysis-tools/sgrna-design). The annealed oligos were cloned into the lentiCRISPR V2 plasmid and used to generate virus. In brief, 1×10^6^ HEK293T cells were plated per well in a 6-well plate. Next day, cells were transfected by mixing 1.1 μg of each sgRNA with 2.7 μg psPAX2, 1 μg pMD2.G and 21 μl TurboFect (Thermo Fisher Scientific, R0531) in 200 μl OPTIMEM (Thermo Fisher Scientific, 31985-062) and incubating for 30 minutes at room temperature (RT) before adding it onto cells. After 24 hours, supernatant was collected and stored at −80 °C while 2 ml fresh media was re-added per well and left for an additional 24 hours. In the meantime, recipient cells were plated at 1×10^5^ cells per well in a 6-well plate. Next day, supernatant was collected from transfected cells, mixed with virus collected the day before and centrifuged for 5 minutes at 1,200 rpm followed by filter sterilisation. To infect recipient cells, 500 μl of virus was mixed with 1.5 ml DMEM and polybrene (8 μg/ml; Sigma-Aldrich, H9268-5G). To generate the double CYRI1/FAM49A KO cells, 250 μl of each virus pool were mixed and used for cell transduction. As a control, parental cells were infected with virus generated from an empty LentiCRISPR V2 plasmid (control KO). After 48 hours post infection, cells were re-plated in selection with 2 μg/ml puromycin (Roth, 0240.3). Antibiotic selection was withdrawn during experiments.

### GST pulldown

Rosetta competent cells were used to grow and purify the respective GST-tagged proteins. GST or GST-RAC1 immobilised on Glutathione Sepharose 4B (GE Healthcare, 17-0756-01) were washed using GST lysis buffer [25 mM Tris-HCL pH 7.2, 150 mM NaCl, 5mM MgCl_2_, 1 % Nonidet P40 (v/v), 5% glycerol (v/v), 1 % protease inhibitor cocktail tablet (Roche, 4693132001), 1 % phosphatase inhibitor cocktails 1 and 2 (v/v; Sigma Aldrich, P5726-5ml and P0044-5ml) in dH_2_O] and used immediately (unloaded) or subjected to nucleotide chelating using 0.5 M EDTA (10 mM final concentration). GST and GST-RAC1 beads were then incubated with Guanosine 5’-diphosphate sodium salt (GDP; 100 μM final concentration; Sigma-Aldrich, G7127) or Guanosine 5’-O-(3-thiotriphosphate) tetralithium salt (GTPγS; 1 mM final concentration; Sigma-Aldrich, G8634) for 15 minutes at 30 °C. Termination of the reaction was achieved by adding MgCl2 (60 mM final concentration). Cells were lysed 24 hours post dox-treatment (1 μg/ml) or transfection in GST lysis buffer and equal protein amounts were incubated with GDP or GTPγS-loaded GST and GST-RAC1 beads for 2 – 72 hours at 4 °C. For GST and GST-CYRI1 pulldown experiments, HEK293T cells were treated with GDP or GTPγS as detailed above prior to incubation with GST and GST-CYRI1 beads overnight (O/N) at 4 °C. Beads were washed following incubation and prepared for western blotting. GST and GST-RAC1 were a kind gift from Angeliki Malliri (Cell Signalling Group, Cancer Research UK Manchester Institute).

### Immunofluorescence

Cells grown on 96-well plates (Greiner Bio, 65589) were fixed in 4 % paraformaldehyde (PFA; ChemCruz, SC281692) for 15 minutes at RT and permeabilized by incubating for 3 minutes with 0.5 % Triton-X (v/v) in Phosphate Buffered Saline (PBS; Thermo Fisher Scientific, 14190-169) at RT. Cells were next blocked in 1 % Bovine Serum Albumin (BSA; Roth, T844.3) in PBS for 30 minutes prior to incubation with relevant primary antibodies (in 1 % BSA) for 1 hour at RT. The respective HRP-conjugated secondary antibodies were then added for an additional hour at RT in the dark. Immunofluorescence was performed using antibodies listed in Supplementary Methods Table 1, as outlined therein. Images were captured using the Yokogawa CQ1 confocal quantitative image cytometer platform (x63 magnification).

### Duolink *in situ* Proximity Ligation Assay (PLA)

Cells were plated in 96-well plates 24 hours prior to fixing in 4 % PFA and permeabilization. For analysis of endogenous CYRI1-RAC1 binding, cells were treated with mouse anti-RAC1 antibody (BD Biosciences, 610650) and rabbit anti-FAM49B antibody (Proteintech, 20127-I-AP). For exogenous interaction the mouse anti-RAC1 antibody was used together with rabbit anti-HA (Covance, PRB-101P). Following incubation with the outlined antibody sets or anti-CYRI1 or anti-HA alone as a control, the respective PLA probes (Olink Bioscience, anti-mouse 92004-0100, anti-rabbit 92002-0100) and the detection reagent kit (Olink Bioscience, 92014-0100) were applied according to manufacturer’s instructions. Alexa fluor 647 Phalloidin and NucBlue Live ReadyProbes Reagent were used as fluorescence markers against the actin cytoskeleton and the nucleus, respectively. Images were taken using the Yokogawa CQ1 confocal quantitative image cytometer platform (x63 magnification). For quantification of duolink count per cell, images were analysed using the built-in CQ1 image analysis software, which was adjusted to count the number of duolink dots and nuclei per image. To account for background, duolink counts per cell from control samples subjected to one antibody only were plotted together with the average duolink count per cell identified for each condition.

### Analysis of cell morphology

Cells were seeded in 6-well plates for 24 hours at varying cellular densities and phase-contrast images were taken on a Zeiss Axiovert 3 (×32 magnification). Cells were classified into normal or pancake-like according to their morphology. By examining parental cells and determining the morphology displayed by the majority of HeLa cells under resting conditions, we classified the normal morphology as cells with an elongated mesenchymal-like phenotype. In contrast, as previously described^31^, cells displaying a flattened morphology with unpolarised membrane ruffling were classified as pancake-like. Images were imported to the ImageJ software and analysed to determine surface area and circularity indices.

### PAK-CRIB pulldown

To detect levels of active RAC1, cells were lysed in GST lysis buffer containing 8 μg/ml biotinylated PAK-CRIB purified peptide and incubated for 30 minutes at 4 °C. Cleared lysates were then incubated with 50 μl Strep-Tactin superflow resin (IBA GmbH, 2-1206-10) for 15 minutes at 4 °C. Beads were washed following incubation and prepared for western blotting to determine levels of active and total RAC1. The biotinylated PAK-CRIB was a kind gift from Angeliki Malliri (Cell Signalling Group, Cancer Research UK Manchester Institute).

### Biochemical analysis of actin polymerisation

Monomeric actin (G-actin) and filamentous actin (F-actin) were extracted from cell lysates using the G-actin/F-actin *in vivo* assay Biochem Kit (Cytoskeleton, Inc., BK037) according to manufacturer’s instructions. Levels of G-actin and F-actin were determined by western blotting using anti-ACTIN antibody provided with the kit. Western blot scans were imported into imageJ and band intensities were quantified as detailed below. Actin polymerisation was calculated as the ratio of F-actin band intensities to G-actin band intensities. As a control, an F-actin enhancing solution, provided in the kit, was used.

### Western blot analysis

Samples were mixed with the appropriate volume of 2x Laemmli sample buffer (Biorad, 1610737) and resolved on 4-20 % (Biorad, 4561094, 4561095 and 4561096), or 12 % Mini-protein TGX precast protein gels (Biorad, 4561043, 451045 and 4561046) or 12 % self-cast gels. Prestained protein ladder (BioFroxx, 1123YL500) was run alongside the samples for protein size reference. Proteins were transferred onto Immobolin PVDF membranes (Millipore, IPFL00010). Western blotting was performed using antibodies listed in Supplementary Methods Table 1, as outlined therein, and visualised on Fuji medical X-ray super RX (Fuji, 4741008389) using the western blotting luminol reagent (SantaCruz Biotechnology, sc-2048 and sc-2049). Band intensities were quantified using the ImageJ Gel analysis tool. Cells and tissue homogenate derived from mice were extracted using T-PER Tissue Extraction Reagent (Thermo Scientific, 78510) with protease inhibitor cocktail (Sigma Aldrich, P8465) as per manufacturer’s protocol. Protein lysates were quantified by Bradford assay (Biorad, 500-0006) and resuspended in Laemmli buffer prior to western blotting.

### Bone marrow chimeras

5–7 week old mice of both sexes were irradiated with two consecutive doses of 450 rads using the RS2000 X-ray machine. After irradiation, mice were reconstituted by intravenous injection of 1.5 × 10^6^ bone marrow cells in sterile PBS. Mice were kept on sterile tap water containing 2 g/l neomycin sulfate (Bioshop) for 3 weeks. Six weeks following irradiation, mice were tested for chimerism by collecting blood from the saphenous vein and looking at the expression of *Cyri1* wild type or mutant allele by isolating DNA and genotyping with TaqMan.

### Generation of *Cyri1* conditional knockout in myeloid cells

*Cyri1* knockout (KO)-first allele (*Fam49b*^tm1a^) on a C57BL/6N background was converted to a conditional allele (*Fam49b*^tm1c^) via Flp recombination^48^. The C57BL/6N background carries a susceptibility allele at *Slc11a1*, which is permissive for exponential growth of *Salmonella* in cells and tissues. We therefore introduced the *Slc11a1* wild type allele (*Slc11a1^G169^*) in the early steps of breeding. Mice carrying the *Fam49b^tm1c^* allele were crossed to the RCS strain BcA17 (87 % C57BL/6J background and 13 % A/J^49^) to introduce the *Slc11a1* wild type allele (Slc11a1^G169^). The resulting mice were serially backcrossed to mice carrying the *Fam49b*^tm1c^ allele to eliminate the A/J background. In parallel, *Fam49b^tm1c/tm1c^* mice were crossed with Meox-Cre mice, leading to the elimination of the floxed exon 6 and the generation of the *Fam49b^tm1d^* allele. From the resultant offspring, we selected *Fam49b^tm1c/tm1d^* mice to be crossed with LysM-Cre, yielding *Fam49b^+/tm1d^* mice with a Cre recombinase under the control of the LysM myeloid-specific promoter (Tg(Lysm-Cre)*Fam49b^+/tm1d^*). Finally, the Tg(Lysm-Cre)*Fam49b^+/tm1d^* were crossed with B6.*Fam49b^tm1c/tm1c^-Slc11a1^G169/G169^* mice, yielding the experimental myeloid-specific conditional KO mice (*Cyri1^−/flox^*) and control mice (*Cyri1^+/flox^*). All mice carry one copy of the wild type allele at *Slc11a1*. Despite the transfer of a *Slc11a1* resistant allele, the C57BL/6N background is more permissive to *Salmonella* growth as compared to the 129X1 background of the *Ity15* mice.

### Bone marrow-derived cells isolation

Femurs and tibias were collected aseptically. After removing most of the muscle and fat, the epiphyses were cut and bones were placed into modified PCR tubes individually hung by the hinge into a 1.5 ml Eppendorf containing 300 μl of sterile PBS. The bone marrow was flushed by short centrifugation at 5,000 rpm for 10 seconds or alternatively using a 25G needle. Red blood cells were lysed with RBC lysis buffer (Sigma Aldrich, 11814389001) incubating for 4 minutes at RT. Cells were pelleted and resuspended in 1 ml sterile PBS or cRPMI (10% FBS, 1% penicillin-streptomycin) media. To generate Bone Marrow-Derived Macrophages (BMDMs), cells were plated in a 10 cm petri dishes supplemented with 30 % M-CSF every two days for a total of six days at 37 °C and 5 % CO_2_.

### *In vitro Salmonella* infection

Parental, control KO and CYRI1 KO HeLa cells were plated on coverslips in 6-well plates for 24 hours after which cells were infected with *S*. Typhimurium (strain SL1344) at a multiplicity of infection (MOI) of 8. After addition of bacteria, cells were incubated for 30 minutes at 37 °C and 5 % CO_2_, before addition of 100 μg/ml gentamicin to kill extracellular bacteria. To assess bacterial entry, cells were washed twice with PBS and prepared for immunofluorescence as outlined above. To visualise intracellular bacteria anti-*Salmonella* antibody (CSA-1; KPL 01-91-99-MG) was utilised. Images were captured using confocal microscopy and analysed in Cell profiler 2.2.0 (rev 9969f42) to calculate the number of bacteria per cell.

### *Salmonella* phagocytosis and replication assays

*Ex vivo* phagocytosis and replication in BMDMs were assessed using an adapted gentamicin protection assay protocol^50^. BMDMs were isolated and cultured as described above. BMDMs were seeded in a 24-well plate on 12 mm diameter glass coverslips at 2–2.5×10^5^ cells per well for 16–24 hours before use. Cells were infected with opsonized *S*. Typhimurium at an MOI of 10–20. After addition of *Salmonella*, plates were centrifuged at 1,400 rpm for 1 minute and incubated for 20 minutes at 37 °C and 5 % CO_2_ before addition of 100 μg/ml gentamicin to kill extracellular bacteria. To assess *Salmonella* phagocytosis, after 30 minutes cells were washed twice with warm PBS and fixed with 2.5 % PFA for 10-30 minutes at 37 °C and 5 % CO_2_. Fixed cells were rinsed three times with PBS and permeabilized with 0.2 % saponin in 10 % normal goat serum O/N at 4 °C. To detect internalised bacteria unconjugated rabbit anti-*Salmonella* antibody (Cedar Lane, B65701R) was used. Images were obtained on a Zeiss LSM 710 confocal microscope and analysed using Zen software and ImageJ.

### Neutrophil enrichment, labelling and 3D migration assay

Total bone marrow neutrophils (~ 80-85 %, Ly6G^+^CD11b^+^) were purified by negative selection using the EasySep Mouse Neutrophil Enrichment kit (StemCell Technologies, 19762) and dye labelled with 10 μM CellTracker Orange (CMTMR; Invitrogen, C2927) in cRPMI for 15 minutes at 37 °C. Cells were suspended in PureCol (Advanced Biomatrix, 5005) and cast in custom built migration chambers at a final collagen concentration of 1.6 mg/ml for 40 minutes, as described previously^51^. Final cell concentrations in the assay were 1–2 × 10^6^ cells per ml gel. Cells were imaged using a Zeiss 880 upright confocal fluorescence microscope for 1 hour at 37 °C. Cells were tracked using the spots tool in Imaris (Bitplane), and speed and directionality parameters were calculated using a custom coded MATLAB program (MathWorks).

### Statistical analysis

Statistical analysis was performed on GraphPad prism. p-values were classified as follows: non-significant corresponds to ns= p > 0.05, while level of significance was denoted as: *= p ≤ 0.05, **= p ≤ 0.01, ***= p ≤ 0.001 and ****= p ≤ 0.0001. Tests are specified in figure legends where appropriate.

### Code availability

All commercial and custom codes used to generate data presented in this study are available from the corresponding authors upon reasonable request.

## Data availability

The data that support the findings of this study are available from the corresponding authors upon reasonable request.

**Supplementary Information** is available in the online version of the paper

## Acknowledgements

We thank Line Larivière, Etienne Flamant, Leïla Rached-D’Astous and Masuda Sader for technical assistance and Nadia Prud’homme and Patricia D’Arcy for mouse breeding and screening. We thank Angeliki Malliri for reagents. We also acknowledge Laura Machesky and Robert Insall for discussions, Francesco Pampaloni for assistance and Kerstin Koch for reviewing the manuscript. This work was supported by the Canadian Institutes of Health Research MOP133700 (to D.M) and by the DFG-funded Collaborative Research Centre on Selective Autophagy (SFB 1177), the European Research Council (ERC) under the European Union’s Horizon 2020 research and innovation program (grant agreement No 742720), the DFG-funded Cluster of Excellence ‘‘Macromolecular Complexes’’ (EXC115), the DFG-funded SPP 1580 program ‘‘Intracellular Compartments as Places of Pathogen-Host-Interactions’’ and by the LOEWE program Ubiquitin Networks (Ub-Net) and the LOEWE Centre for Gene and Cell Therapy Frankfurt (CGT), which are both funded by the State of Hessen, Germany (to I.D). Funding had been provided to M.M.E. by Le Fonds de Recherche Santé du Québec (FRQS) and to H.M. by the Alexander von Humboldt foundation as a Humboldt research fellowship for postdoctoral researchers.

## Author Contributions

K.E.Y., H.M., E.F., and M.M.E contributed equally to this work. K.E.Y., H.M., E.F., M.M.E., S.M.V., MC, D.M. and I.D. designed the conceptual framework of the study and experiments; K.E.Y., and M.M.E. contributed to ENU mutation identification and performed all *in vivo* and *ex vivo* experiments involving *Cyri1* mutant and conditional alleles and analysed data; H.M. designed, performed and analysed all biochemical and cell biology assays; E.F. designed and performed *in vitro Salmonella* infection experiments and performed some biochemical assays; J.A.S. and J.M. performed analyses of exome sequences; A.G. and J.N.M. designed, performed and analysed experiments with neutrophil mobility assays; D.M. and I.D. supervised the project. All authors wrote the manuscript.

## Author Information

The authors declare no competing financial interests. Correspondence and requests for materials should be addressed to I.D. or D.M.

## References

1 Kirk, M. D. et al. World Health Organization Estimates of the Global and Regional Disease Burden of 22 Foodborne Bacterial, Protozoal, and Viral Diseases, 2010: A Data Synthesis. PLoS Med 12, e1001921, doi: 10.1371/journal.pmed.1001921 (2015).

2 Majowicz, S. E. et al. The global burden of nontyphoidal Salmonella gastroenteritis. Clin Infect Dis 50, 882–889, doi:10.1086/650733 (2010).

3 Prestinaci, F., Pezzotti, P. & Pantosti, A. Antimicrobial resistance: a global multifaceted phenomenon. Pathog Glob Health 109, 309–318, doi: 10.1179/2047773215Y.0000000030 (2015).

4 Fabrega, A. & Vila, J. Salmonella enterica serovar Typhimurium skills to succeed in the host: virulence and regulation. Clin Microbiol Rev 26, 308–341, doi:10.1128/CMR.00066-12 (2013).

5 Rydstrom, A. & Wick, M. J. Monocyte recruitment, activation, and function in the gut-associated lymphoid tissue during oral Salmonella infection. J Immunol 178, 5789–5801 (2007).

6 Worley, M. J., Nieman, G. S., Geddes, K. & Heffron, F. Salmonella typhimurium disseminates within its host by manipulating the motility of infected cells. Proceedings of the National Academy of Sciences of the United States of America 103, 17915–17920, doi:10.1073/pnas.0604054103 (2006).

7 Ohl, M. E. & Miller, S. I. Salmonella: a model for bacterial pathogenesis. Annu Rev Med 52, 259–274, doi:10.1146/annurev.med.52.1.259 (2001).

8 LaRock, D. L., Chaudhary, A. & Miller, S. I. Salmonellae interactions with host processes. Nat Rev Microbiol 13, 191–205, doi:10.1038/nrmicro3420 (2015).

9 Schlumberger, M. C. & Hardt, W. D. Salmonella type III secretion effectors: pulling the host cell’s strings. Curr Opin Microbiol 9, 46–54, doi:10.1016/j.mib.2005.12.006 (2006).

10 Byndloss, M. X., Rivera-Chavez, F., Tsolis, R. M. & Baumler, A. J. How bacterial pathogens use type III and type IV secretion systems to facilitate their transmission. Curr Opin Microbiol 35, 1–7, doi:10.1016/j.mib.2016.08.007 (2017).

11 Guiney, D. G. & Lesnick, M. Targeting of the actin cytoskeleton during infection by Salmonella strains. Clin Immunol 114, 248–255, doi:10.1016/j.clim.2004.07.014 (2005).

12 Wood, M. W., Rosqvist, R., Mullan, P. B., Edwards, M. H. & Galyov, E. E. SopE, a secreted protein of Salmonella dublin, is translocated into the target eukaryotic cell via a sip-dependent mechanism and promotes bacterial entry. Mol Microbiol 22, 327–338 (1996).

13 Friebel, A. et al. SopE and SopE2 from Salmonella typhimurium activate different sets of RhoGTPases of the host cell. J Biol Chem 276, 34035–34040, doi: 10.1074/jbc.M100609200 (2001).

14 Bakshi, C. S. et al. Identification of SopE2, a Salmonella secreted protein which is highly homologous to SopE and involved in bacterial invasion of epithelial cells. J Bacteriol 182, 2341–2344 (2000).

15 Hardt, W. D., Chen, L. M., Schuebel, K. E., Bustelo, X. R. & Galan, J. E. S. typhimurium encodes an activator of Rho GTPases that induces membrane ruffling and nuclear responses in host cells. Cell 93, 815–826 (1998).

16 Orchard, R. C. & Alto, N. M. Mimicking GEFs: a common theme for bacterial pathogens. Cell Microbiol 14, 10–18, doi:10.1111/j.1462-5822.2011.01703.x (2012).

17 Humphreys, D., Davidson, A., Hume, P. J. & Koronakis, V. Salmonella virulence effector SopE and Host GEF ARNO cooperate to recruit and activate WAVE to trigger bacterial invasion. Cell Host Microbe 11, 129–139, doi:10.1016/j.chom.2012.01.006 (2012).

18 Gautreau, A. et al. Purification and architecture of the ubiquitous Wave complex. Proceedings of the National Academy of Sciences of the United States of America 101, 4379–4383, doi:10.1073/pnas.0400628101 (2004).

19 Eden, S., Rohatgi, R., Podtelejnikov, A. V., Mann, M. & Kirschner, M. W. Mechanism of regulation of WAVE1-induced actin nucleation by Rac1 and Nck. Nature 418, 790–793, doi:10.1038/nature00859 (2002).

20 Ismail, A. M., Padrick, S. B., Chen, B., Umetani, J. & Rosen, M. K. The WAVE regulatory complex is inhibited. Nature structural & molecular biology 16, 561–563, doi: 10.1038/nsmb. 1587 (2009).

21 Chen, Z. et al. Structure and control of the actin regulatory WAVE complex. Nature 468, 533–538, doi:10.1038/nature09623 (2010).

22 Francis, C. L., Ryan, T. A., Jones, B. D., Smith, S. J. & Falkow, S. Ruffles induced by Salmonella and other stimuli direct macropinocytosis of bacteria. Nature 364, 639–642, doi:10.1038/364639a0 (1993).

23 Steele-Mortimer, O. The Salmonella-containing vacuole: moving with the times. Curr Opin Microbiol 11, 38–45, doi:10.1016/j.mib.2008.01.002 (2008).

24 Ham, H., Sreelatha, A. & Orth, K. Manipulation of host membranes by bacterial effectors. Nat Rev Microbiol 9, 635–646, doi:10.1038/nrmicro2602 (2011).

25 Figueira, R. & Holden, D. W. Functions of the Salmonella pathogenicity island 2 (SPI-2) type III secretion system effectors. Microbiology 158, 1147–1161, doi: 10.1099/mic.0.058115-0 (2012).

26 Schleker, S. et al. The current Salmonella-host interactome. Proteomics Clin Appl 6, 117–133, doi:10.1002/prca.201100083 (2012).

27 Mostowy, S. & Shenoy, A. R. The cytoskeleton in cell-autonomous immunity: structural determinants of host defence. Nat Rev Immunol 15, 559–573, doi:10.1038/nri3877 (2015).

28 Hendriksen, R. S. et al. Global monitoring of Salmonella serovar distribution from the World Health Organization Global Foodborne Infections Network Country Data Bank: results of quality assured laboratories from 2001 to 2007. Foodborne Pathog Dis 8, 887–900, doi:10.1089/fpd.2010.0787 (2011).

29 Caignard, G. et al. Mouse ENU Mutagenesis to Understand Immunity to Infection: Methods, Selected Examples, and Perspectives. Genes (Basel) 5, 887–925, doi: 10.3390/genes5040887 (2014).

30 Soding, J., Biegert, A. & Lupas, A. N. The HHpred interactive server for protein homology detection and structure prediction. Nucleic acids research 33, W244–248, doi: 10.1093/nar/gki408 (2005).

31 Michiels, F., Habets, G. G., Stam, J. C., van der Kammen, R. A. & Collard, J. G. A role for Rac in Tiam1-induced membrane ruffling and invasion. Nature 375, 338–340, doi: 10.1038/375338a0 (1995).

32 May, R. C. & Machesky, L. M. Phagocytosis and the actin cytoskeleton. Journal of cell science 114, 1061–1077 (2001).

33 Krause, M. & Gautreau, A. Steering cell migration: lamellipodium dynamics and the regulation of directional persistence. Nature reviews. Molecular cell biology 15, 577–590, doi:10.1038/nrm3861 (2014).

34 Wijburg, O. L., Simmons, C. P., van Rooijen, N. & Strugnell, R. A. Dual role for macrophages in vivo in pathogenesis and control of murine Salmonella enterica var. Typhimurium infections. Eur J Immunol 30, 944–953, doi:10.1002/1521-4141 (200003)30:3<944::AID-IMMU944>3.0.CO;2-1 (2000).

35 Richter-Dahlfors, A., Buchan, A. M. & Finlay, B. B. Murine salmonellosis studied by confocal microscopy: Salmonella typhimurium resides intracellularly inside macrophages and exerts a cytotoxic effect on phagocytes in vivo. The Journal of experimental medicine 186, 569–580 (1997).

36 Campa, C. C. et al. Rac signal adaptation controls neutrophil mobilization from the bone marrow. Sci Signal 9, ra124, doi:10.1126/scisignal.aah5882 (2016).

37 Pankov, R. et al. A Rac switch regulates random versus directionally persistent cell migration. The Journal of cell biology 170, 793–802, doi: 10.1083/jcb.200503152 (2005).

38 Finlay, B. B. & McFadden, G. Anti-immunology: evasion of the host immune system by bacterial and viral pathogens. Cell 124, 767–782, doi: 10.1016/j.cell.2006.01.034 (2006).

39 Broz, P., Ohlson, M. B. & Monack, D. M. Innate immune response to Salmonella typhimurium, a model enteric pathogen. Gut Microbes 3, 62–70, doi:10.4161/gmic.19141 (2012).

40 Uhlen, M. et al. Proteomics. Tissue-based map of the human proteome. Science 347, 1260419, doi: 10.1126/science.1260419 (2015).

41 Marei, H. & Malliri, A. Rac1 in human diseases: The therapeutic potential of targeting Rac1 signaling regulatory mechanisms. Small GTPases 8, 139–163, doi: 10.1080/21541248.2016.1211398 (2017).

42 Johansson, C., Ingman, M. & Jo Wick, M. Elevated neutrophil, macrophage and dendritic cell numbers characterize immune cell populations in mice chronically infected with Salmonella. Microb Pathog 41, 49–58, doi: 10.1016/j.micpath.2006.03.004 (2006).

43 Nedelec, Y. et al. Genetic Ancestry and Natural Selection Drive Population Differences in Immune Responses to Pathogens. Cell 167, 657–669 e621, doi:10.1016/j.cell.2016.09.025 (2016).

44 Fiskin, E., Bionda, T., Dikic, I. & Behrends, C. Global Analysis of Host and Bacterial Ubiquitinome in Response to Salmonella Typhimurium Infection. Mol Cell 62, 967–981, doi:10.1016/j.molcel.2016.04.015 (2016).

45 Roy, M. F. et al. Pyruvate kinase deficiency confers susceptibility to Salmonella typhimurium infection in mice. The Journal of experimental medicine 204, 2949–2961, doi:10.1084/jem.20062606 (2007).

46 Eva, M. M. et al. Altered IFN-gamma-mediated immunity and transcriptional expression patterns in N-Ethyl-N-nitrosourea-induced STAT4 mutants confer susceptibility to acute typhoid-like disease. J Immunol 192, 259–270, doi: 10.4049/jimmunol. 1301370 (2014).

47 Marei, H. et al. Differential Rac1 signalling by guanine nucleotide exchange factors implicates FLII in regulating Rac1-driven cell migration. Nat Commun 7, 10664, doi:10.1038/ncomms10664 (2016).

48 Skarnes, W. C. et al. A conditional knockout resource for the genome-wide study of mouse gene function. Nature 474, 337–342, doi:10.1038/nature10163 (2011).

49 Fortin, A. et al. Recombinant congenic strains derived from A/J and C57BL/6J: a tool for genetic dissection of complex traits. Genomics 74, 21–35, doi: 10.1006/geno.2001.6528 (2001).

50 Cunnington, A. J., de Souza, J. B., Walther, M. & Riley, E. M. Malaria impairs resistance to Salmonella through heme- and heme oxygenase-dependent dysfunctional granulocyte mobilization. Nature medicine 18, 120–127, doi: 10.1038/nm.2601 (2011).

51 Sixt, M. & Lammermann, T. In vitro analysis of chemotactic leukocyte migration in 3D environments. Methods in molecular biology 769, 149–165, doi:10.1007/978-1-61779-207-6_11 (2011).

